# Hypusinated eIF5A is expressed in pancreas and spleen of individuals with type 1 and type 2 diabetes

**DOI:** 10.1101/745919

**Authors:** Teresa L. Mastracci, Stephanie C. Colvin, Leah R. Padgett, Raghavendra G. Mirmira

## Abstract

Eukaryotic initiation factor 5A (*EIF5A*) is found in diabetes-susceptibility loci in mouse and human. eIF5A is the only protein known to contain hypusine (<u>hy</u>droxy<u>pu</u>trescine ly<u>sine</u>), a polyamine-derived amino acid formed post-translationally in a reaction catalyzed by deoxyhypusine synthase (DHPS). Previous studies showed pharmacologic blockade of DHPS in type 1 diabetic NOD mice and type 2 diabetic db/db mice improved glucose tolerance and preserved beta-cell mass, which suggests that hypusinated eIF5A (eIF5A^Hyp^) may play a role in diabetes pathogenesis by direct action on the beta cells and/or altering the adaptive or innate immune responses. To translate these findings to human, we examined tissue from individuals with and without type 1 and type 2 diabetes to determine the expression of eIF5A^Hyp^. We detected eIF5A^Hyp^ in beta cells, exocrine cells and immune cells; however, there was also unexpected enrichment of eIF5A^Hyp^ in pancreatic polypeptide-expressing PP cells. Interestingly, the presence of eIF5A^Hyp^ co-expressing PP cells was not enhanced with disease. These data identify new aspects of eIF5A biology and highlight the need to examine human tissue to understand disease.

## INTRODUCTION

The mechanisms underlying the pathogeneses of type 1 diabetes (T1D) and type 2 diabetes (T2D) involve the activation of systemic and local inflammatory pathways, leading to eventual dysfunction, de-differentiation and/or death of the beta cells in the pancreatic islet. Elucidating the molecular mechanisms driving the inflammatory response is applicable to the development of therapies for both diseases. In addition, an urgent priority in T1D research is the discovery of biomarkers that can assist in the identification of individuals with pre-clinical disease so early preventative therapeutic interventions can be implemented.

Recently, our laboratories have been investigating the involvement of the hypusinated form of eukaryotic initiation factor 5A (eIF5A) in the development and progression of diabetes in mice. To date, eIF5A is the only known protein to contain hypusine (<u>hy</u>droxy<u>pu</u>trescine ly<u>sine</u>) [1], which is a polyamine-derived amino acid. This post-translational modification, formed by the process of “hypusination” [2], is catalyzed through a multi-step reaction initiated by the *rate-limiting* enzyme deoxyhypusine synthase (DHPS) and uses the polyamine spermidine as a cofactor to modify the Lys50 of eIF5A [2]. Previous studies using human cell lines and yeast determined that eIF5A, the hypusinated form of eIF5A (eIF5A^Hyp^) and DHPS are vital for cell viability and proliferation [3,4]. Evolutionarily, eIF5A is highly conserved including the amino acid sequence surrounding the hypusine residue, which suggests an important role for this modification [5]. Whereas studies across species have established that eIF5A^Hyp^ efficiently binds the ribosome complex and facilitates mRNA translation [3,6,7], the exact function of eIF5A and eIF5A^Hyp^ remains unknown..

Interestingly, the gene encoding eIF5A is found in the *Idd4* diabetes-susceptibility locus in non-obese diabetes (NOD) mice [8,9]. In prior studies, we showed that eIF5A^Hyp^ is responsible for the translation of a subset of cytokine-induced transcripts in beta cells in mouse models of diabetes [10,11], and that eIF5A^Hyp^ also appears to be required for the activation and proliferation of effector T helper cells [12]. Moreover, reducing the hypusination of eIF5A in NOD mice, a model of T1D, by pharmacological inhibition of DHPS resulted in reduced insulitis, improved glucose tolerance and preserved beta cell mass [12]. Similarly, pharmacological blockade of DHPS in db/db mice [13], a model of T2D improved glucose tolerance and enhanced beta cell mass [14]. Together these data suggest that eIF5A^Hyp^ may play a critical role in the pathogenesis of diabetes and altering the expression of eIF5A^Hyp^ may improve diabetes outcomes long-term.

To translate these findings to human, a greater understanding of eIF5A^Hyp^ in the human pancreas and spleen would be required. In particular, determining the expression pattern of eIF5A^Hyp^ in human and whether eIF5A^Hyp^-expressing cells stratify with characteristics of disease would be informative. In this study, we used human donor tissue samples from the Network of Pancreatic Organ Donors with Diabetes (nPOD) (www.jdrfnpod.org) to examine the cell-type distribution of eIF5A^Hyp^ in the human pancreas and spleen from individuals with T1D, T2D and non-diabetic controls.

## RESULTS

### Beta cell and non-beta cell distribution of eIF5A^Hyp^ in mouse

We previously developed and characterized a novel antibody that recognizes the unique amino acid hypusine, formed exclusively through posttranslational modification of the Lys50 residue of eIF5A (eIF5A^Hyp^) [10]. In this study, we utilized this antibody to investigate the expression of eIF5A^Hyp^ in mouse and human pancreas tissue and isolated islets as well as human spleen tissue, to characterize the expression pattern of eIF5A^Hyp^ and determine if eIF5A^Hyp^-expressing cells stratify with characteristics of disease. To that end, we first confirmed the presence of eIF5A^Hyp^ in islets isolated from mouse and human pancreas as well as in mouse pancreas and human acinar (exocrine) tissue (Fig 1A).

**Figure 1.**
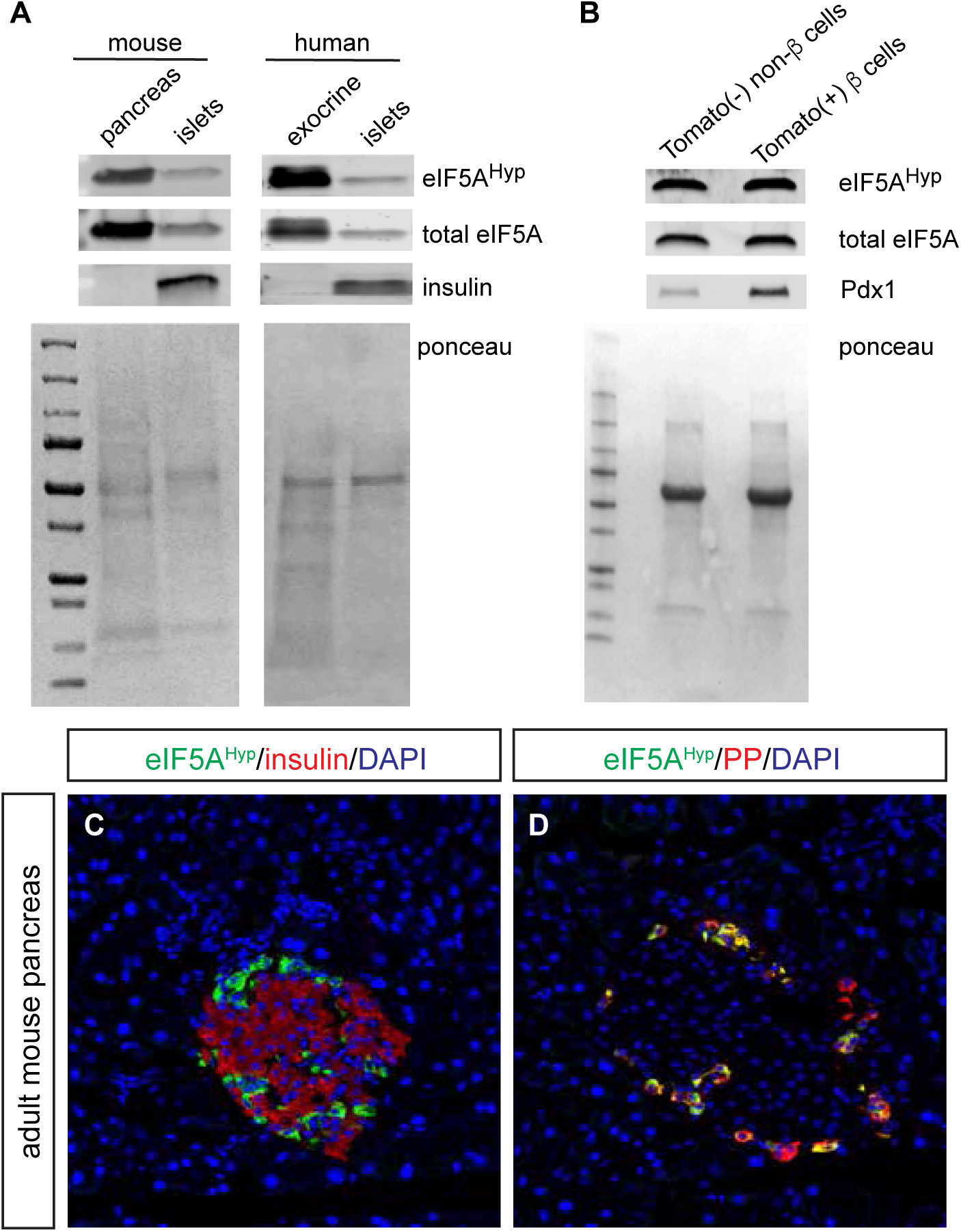
Expression of hypusinated eIF5A (eIF5A^Hyp^) in mouse and human pancreatic islets. A Western blot from mouse and human pancreas tissue and isolated pancreatic islets. B Western blot from FACS sorted mouse islet cell populations. C, D Representative immunofluorescence images of mouse tissue demonstrating robust expression of eIF5A^Hyp^ in PP-expressing cells.

We next utilized the *RIP-cre;R26R*^*Tomato*^ mouse model wherein the insulin-expressing cells were labeled with a lineage trace, thereby generating beta cells indelibly marked with fluorescent reporter (tomato) expression. Islet cells from *RIP-cre;R26R*^*Tomato*^ and control animals were sorted by FACS, using the presence and absence of tomato expression to separate cells into two populations: beta cells (tomato-positive) and non-beta cells (tomato-negative). The cell types represented in the “non-beta cell” sample would include (ordered from largest population to smallest): glucagon-expressing alpha cells, somatostatin-expressing delta cells, pancreatic polypeptide-expressing PP cells, ghrelin-expressing epsilon cells, exocrine cells (a possible contaminant from the process of islet isolation) and support cells including endothelial cells. A similar quantity of tomato-positive beta cells (1.92×10^5^ cells) and tomato-negative non-beta cells (2.13×10^5^ cells) were collected (Fig EV1). Subsequent western blot analysis identified that eIF5A^Hyp^ was present in nearly identical abundance in both the beta cell (tomato-positive) and non-beta cell (tomato-negative) populations (Fig 1B). The expression of Pdx1 confirms the enrichment of beta cells in the tomato positive cells; the lower level of Pdx1 expression in the non-beta cell fraction can be attributed to the somatostatin-expressing delta cells. These data demonstrate that eIF5A^Hyp^ is expressed in both the beta cell and non-beta cell fractions; however, the specific non-beta cell type(s) expressing eIF5A^Hyp^ cannot be clarified from these data. Therefore, to characterize the spatial distribution of eIF5A^Hyp^ expression pattern in the islet, we performed co-immunofluorescence staining for eIF5A^Hyp^ and islet hormones in mouse pancreas tissue. Whereas relatively weak immunostaining of eIF5A^Hyp^ was found throughout the pancreas and islets, robust immunostaining of eIF5A^Hyp^ was found in the islet cell population that expressed pancreatic polypeptide (Fig 1C, D). Based on the known abundance of islet cell populations [15–17], PP cells would represent only a small proportion of the “tomato-negative non-beta cell” FACS sample, yet the abundance of eIF5A^Hyp^ expression is nearly equivalent to that observed in the pure “tomato-positive beta cell” FACS sample (Fig 1B). Thus, the robust expression of eIF5A^Hyp^ in the PP cells is consistent with the immunoblot data.

### eIF5A^Hyp^-expressing cells in the pancreas of human type 2 diabetes

To characterize the expression pattern of eIF5A^Hyp^ in the human pancreas, we utilized tissue samples from the Network of Pancreatic Organ Donors with Diabetes (nPOD). A cohort of tissues from donors with and without T2D were provided (Table 1). Both pancreas and spleen tissues were acquired from each donor; age, gender, ethnicity and BMI were matched where possible. Given the relatively small size of the cohort, quantitative evaluations were not possible. Therefore, we evaluated the presence or absence of eIF5A^Hyp^, its cell-type expression pattern, and its expression correlation with disease.

**Table 1.** Human donor pancreas and spleen tissue from T2D and matched controls.

| nPOD case # | Sample name | Age (years) | Gender (male/female) | Ethnicity | BMI | T2D (yes/no) | C-peptide (ng/mL) | HbA1c |
| --- | --- | --- | --- | --- | --- | --- | --- | --- |
| nPOD-6097 | F1-control | 43.1 | female | Caucasian | 36.4 | no | 16.76 | 7.1 |
| nPOD-6102 | F2-control | 45.1 | female | Caucasian | 35.1 | no | 0.55 | 6.1 |
| nPOD-6132 | F1-T2D | 55.8 | female | Hispanic | 44.6 | yes | 0.8 | 9.1 |
| nPOD-6109 | F2-T2D | 48.8 | female | Hispanic | 32.5 | yes | <0.05 | 8 |
| nPOD-6060 | M1-control | 24 | male | Caucasian | 32.7 | no | 13.63 | N/A |
| nPOD-6091 | M2-control | 27.1 | male | Caucasian | 35.6 | no | 7.71 | 6.3 |
| nPOD-6114 | M1-T2D | 42.8 | male | Caucasian | 31 | yes | 0.58 | 7.8 |
| nPOD-6188 | M2-T2D | 36.1 | male | Hispanic | 30.6 | yes | 3.45 | 7.2 |

Pancreas tissue sections were co-immunostained with the eIF5A^Hyp^-specific antibody and antibodies that recognized the hormones expressed by each of the endocrine cell populations in the islet (insulin, glucagon, somatostatin, ghrelin and pancreatic polypeptide). Co-localization was not observed between eIF5A^Hyp^ and insulin (Fig 2A,B), glucagon (Fig 2C,D), ghrelin (Fig 2E,F), or somatostatin (Fig 2G,H). However, as observed in the mouse pancreas, cells expressing pancreatic polypeptide were identified to co-express high levels of eIF5A^Hyp^ in control pancreas tissue (Fig 3A). These cells also expressed chromograninA, which confirms their identity as neuroendocrine cells (Fig 3B). The co-localization of eIF5A^Hyp^ with pancreatic polypeptide in the PP-expressing cells was observed in pancreas tissues from donors with T2D (Fig 3C,D) and non-diabetic controls, suggesting no stratification with disease. Notably, whereas PP and eIF5A^Hyp^ were expressed in the same cells, the expression pattern reveals localization in different compartments, with the eIF5A^Hyp^ pattern suggestive of localization in the endoplasmic reticulum (Fig 3E).

**Figure 2.**
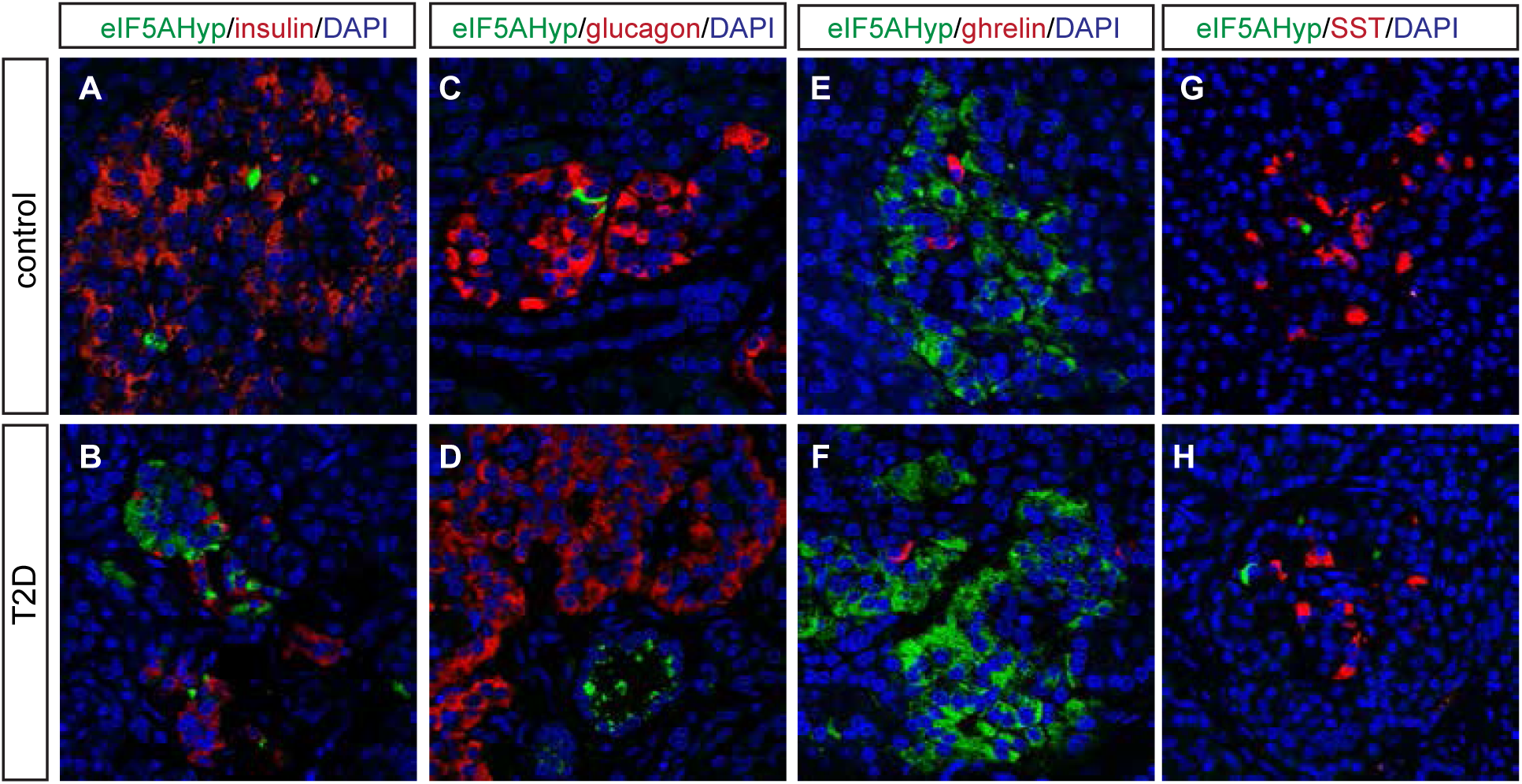
The expression pattern of eIF5A^Hyp^ in T2D and control pancreatic tissue. In controls (matched for age, gender and BMI) and T2D pancreas, we evaluated the co-expression of eIF5A^Hyp^ with all islet hormones and found no overlap with insulin (A,B), glucagon (C,D), ghrelin (E,F) or somatostatin (G,H).

**Figure 3.**
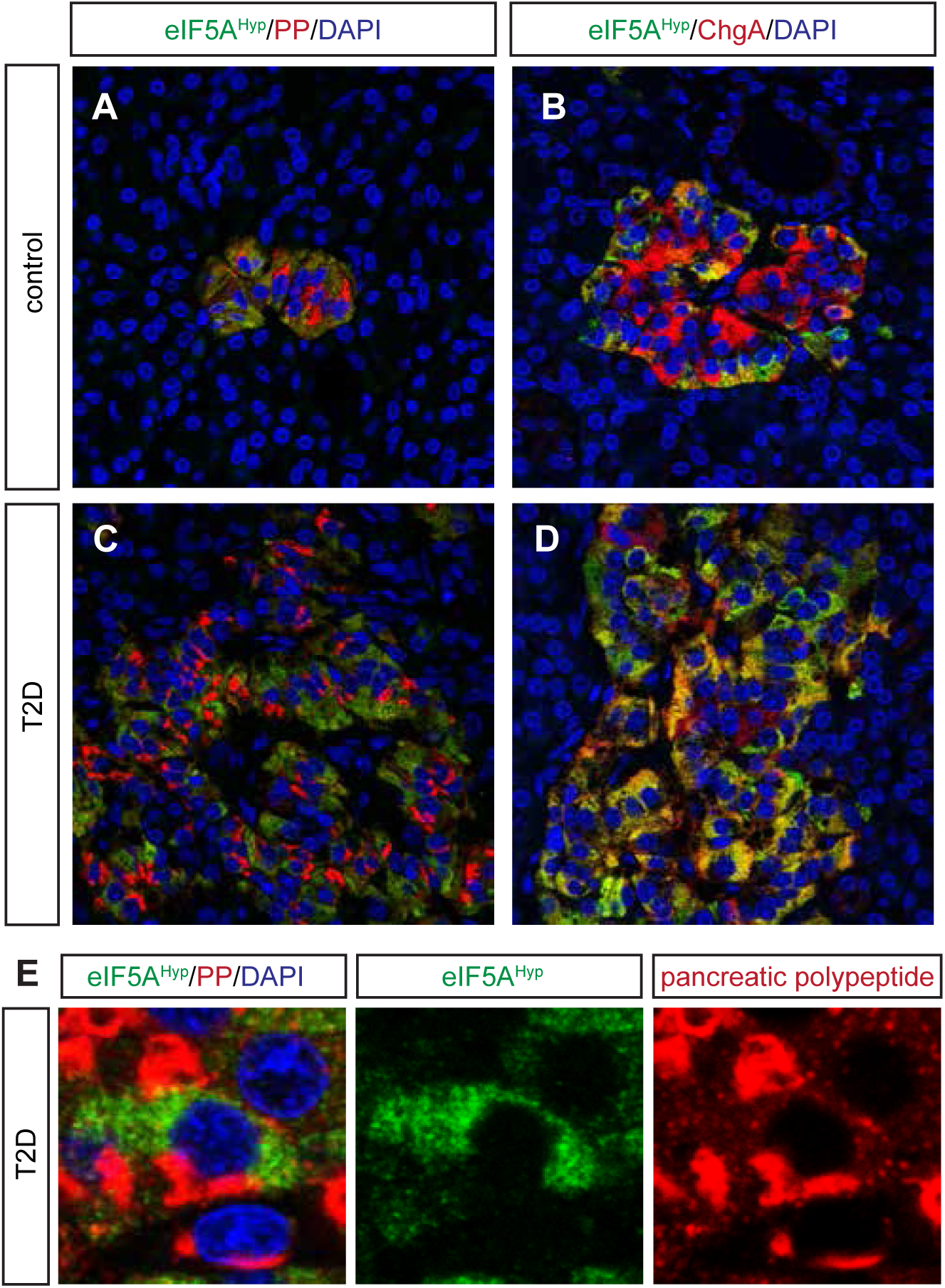
eIF5A^Hyp^ is robustly expressed in the pancreatic polypeptide-expressing PP cells in the islet. A-D In both controls and T2D pancreas, co-expression of eIF5A^Hyp^ with pancreatic polypeptide (PP) and chromograninA (ChgA) was observed. E An expression pattern of eIF5A^Hyp^ in the PP cells suggestive of localization to the ER was observed in cells in both controls and T2D pancreas.

Spleen tissue sections from the same donors were co-immunostained with eIF5A^Hyp^ and markers of various cell types. In particular, Pax5-expressing B cells, CD4-expressing T cells, and CD8-expressing T cells were evaluated for co-expression of eIF5A^Hyp^. Whereas most Pax5+ B cells expressed eIF5A^Hyp^, a select group of eIF5A^Hyp^-expressing cells co-expressed either CD4 or CD8 (Fig 4A-C). No differences in staining intensity or distribution were observed between samples from T2D and controls (Fig 4D-E).

**Figure 4.**
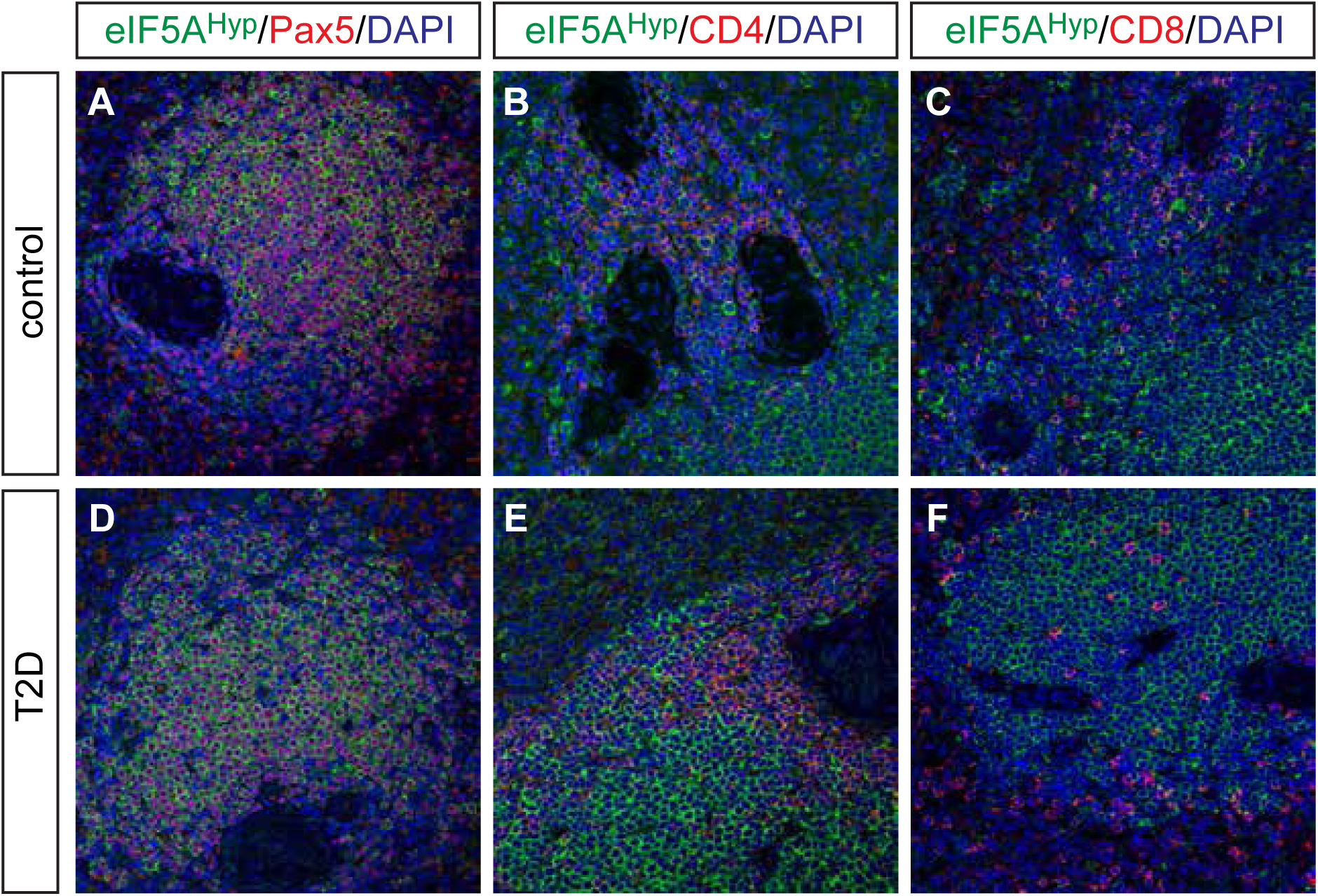
eIF5A^Hyp^ expression pattern in the spleen of control and T2D. eIF5A^Hyp^ is expressed in immune cells in the spleen. We evaluated expression of eIF5A^Hyp^ in Pax5+ B cells, CD4+ T cells and CD8+ T cells in the spleens of donors with T2D and controls matched for age, gender and BMI. Most eIF5A^Hyp^+ cells co-expressed Pax5+ (A,D); however, a select group of eIF5A^Hyp^+ cells expressed either CD4+ (B,E) or CD8+ (C,F).

### eIF5A^Hyp^-expressing cells in the pancreas of human type 1 diabetes

Donor pancreas and spleen tissue from individuals with T1D were also acquired from nPOD and evaluated for eIF5A^Hyp^ expression. This cohort of samples included T1D donors that were autoantibody-positive and autoantibody-negative, with both short and long disease duration; non-diabetic controls were matched for age, gender, ethnicity and BMI (Table 2). Similar to the T2D/control samples, we identified cells co-expressing the hormone PP with high intensity eIF5A^Hyp^ immunostaining (Fig 5A-F); co-expression of eIF5A^Hyp^ with other islet hormones was not observed. Moreover, the eIF5A^Hyp^-expressing cells expressed ChromograninA (Fig 5 G-I), which again confirmed that these cells are neuroendocrine in nature. Evaluation of spleen tissue for all T1D donors and controls revealed an identical pattern of expression to that observed in the T2D donors and controls. Specifically, the majority of eIF5A^Hyp^-expressing cells co-expressed Pax5 (Fig 6).

**Table 2.** Human donor pancreas and spleen tissue from T1D and matched controls.

| nPOD case # | Sample name | Age (years) | Gender (male/female) | Ethnicity | BMI | T1D (yes/no) | c-peptide | Auto Antibodies Detected | Duration of disease (years) |
| --- | --- | --- | --- | --- | --- | --- | --- | --- | --- |
| nPOD-6179 | 1-control | 21.8 | female | Caucasian | 20.7 | no | 2.74 | N/A | N/A |
| nPOD-6224 | 1-AAb-negative | 21 | female | Caucasian | 22.8 | yes | <0.05 | negative | 1.5 |
| nPOD-6070 | 1-AAb-positive | 22.6 | female | Caucasian | 21.6 | yes | <0.05 | mIAA+;1A-2A+ | 7 |
| nPOD-6034 | 2-control | 32 | female | Caucasian | 25.2 | no | 3.15 | N/A | N/A |
| nPOD-6121 | 2-AAb-negative | 33.9 | female | Caucasian | 18 | yes | 0.24 | negative | 4 |
| nPOD-6143 | 2-AAb-positive | 32.6 | female | Caucasian | 26.1 | yes | <0.05 | 1A-2A+;mIAA+ | 7 |
| nPOD-6229 | 3-control | 31 | female | Caucasian | 26.9 | no | 6.23 | N/A | N/A |
| nPOD-6208 | 3-AAb-negative | 32 | female | Caucasian | 23.4 | yes | <0.05 | negative | 16 |
| nPOD-6077 | 3-AAb-positive | 32.9 | female | Caucasian | 22 | yes | <0.05 | mIAA+ | 18 |
| nPOD-6015 | 4-control | 39 | female | Caucasian | 32.2 | no | 1.99 | N/A | N/A |
| nPOD-6038 | 4-AAb-negative | 37.2 | female | Caucasian | 30.9 | yes | 0.2 | negative | 20 |
| nPOD-6054 | 4-AAb-positive | 35.1 | female | Caucasian | 30.4 | yes | <0.05 | mIAA+ | 30 |
| nPOD-6055 | 5-control | 27 | male | Caucasian | 22.7 | no | 0.59 | N/A | N/A |
| nPOD-6041 | 5-AAb-negative | 26.3 | male | Caucasian | 28.4 | yes | <0.05 | negative | 10 |
| nPOD-6180 | 5-AAb-positive | 27.1 | male | Caucasian | 25.9 | yes | <0.05 | GADA+;1A-2A+;<br>ZnT8A+;mIAA+ | 11 |
| nPOD-6104 | 6-control | 41 | male | Caucasian | 20.5 | no | 20.55 | N/A | N/A |
| nPOD-6173 | 6-AAb-negative | 44.1 | male | Caucasian | 23.9 | yes | <0.05 | negative | 15 |
| nPOD-6141 | 6-AAb-positive | 36.7 | male | Caucasian | 26 | yes | <0.05 | GADA+;1A-2A+;<br>ZnT8A+;mIAA+ | 28 |

**Figure 5.**
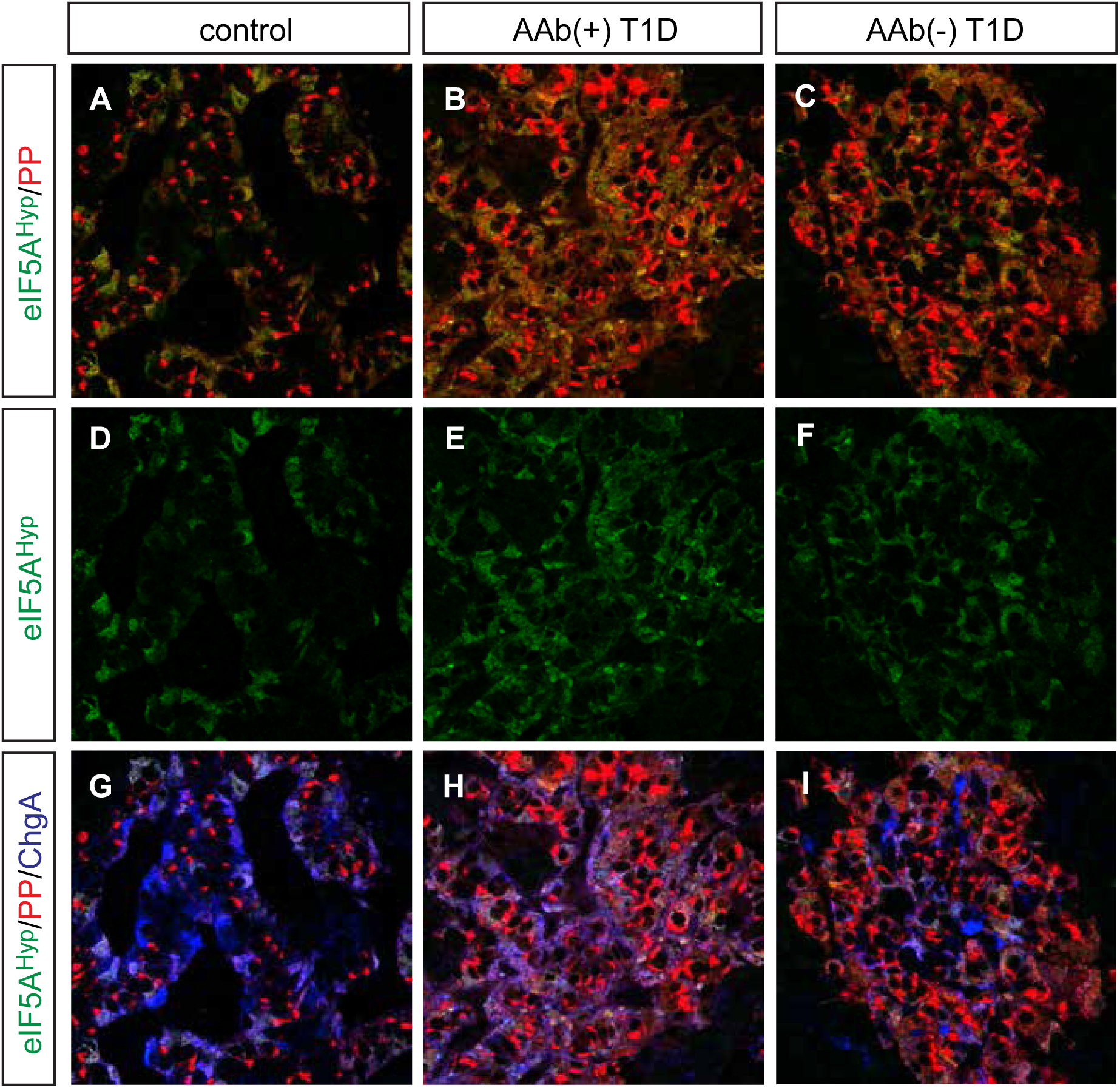
Expression of eIF5A^Hyp^ in the T1D pancreas. A-F Identical to the pattern identified in T2D and control tissues, high expression of eIF5A^Hyp^ is observed in PP cells in the T1D, both auto-antibody positive (aAb+) and auto-antibody negative(aAb-), pancreas and controls (matched for age, gender, ethnicity and BMI). G-I In all cases, these cells express the endocrine cell marker ChromograninA (ChgA).

**Figure 6.**
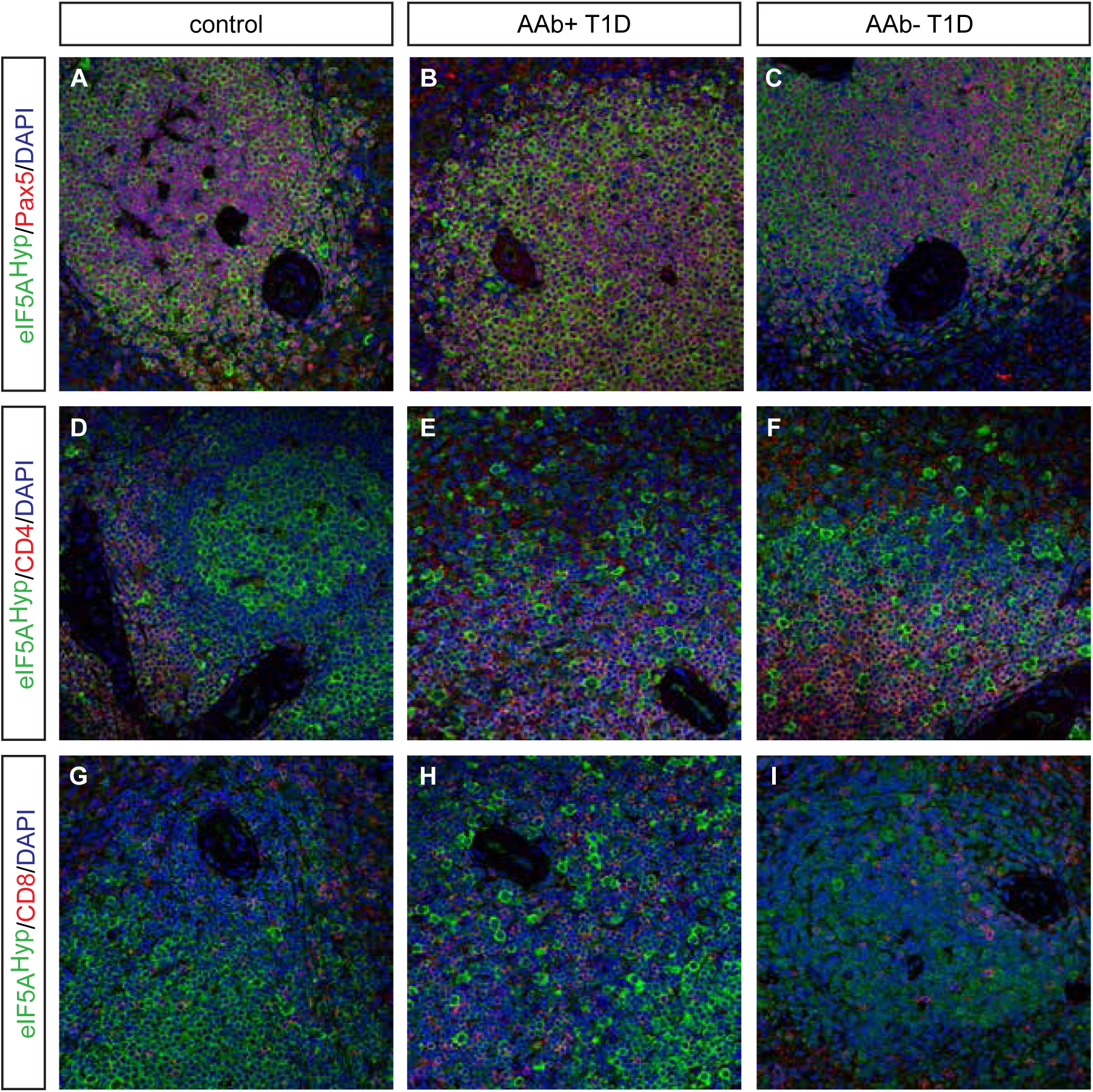
Expression of eIF5A^Hyp^ in spleen tissue from donors with T1D and matched control donors. We examined spleen tissue from persons with autoantibody positive (AAb+) and auto-antibody negative (AAb-) T1D, and corresponding controls matched for age, gender, ethnicity, and BMI. As observed in T2D and matched control spleen tissue, most eIF5A^Hyp^-expressing cells were Pax5+ (A-C); however, some eIF5A^Hyp^+ cells expressed either CD4+ (D-F) or CD8+ (G-I).

## DISCUSSION

Previous data from mouse models identified that pharmacological modulation of the hypusination of eIF5A enhanced beta cell mass and improved glucose tolerance in mouse models of both T1D and T2D [12,14], thereby suggesting an important role for eIF5A^Hyp^ in the setting of diabetes. However, to translate these findings to human, a greater understanding of eIF5A^Hyp^ in the human pancreas and spleen would be required. This study represents the first description of eIF5A^Hyp^ expression in human organs from donors with and without diabetes. Importantly, our results reveal a heretofore unappreciated cell-specific enrichment of eIF5A^Hyp^ in subsets of endocrine cells in the pancreas and immune cells in the spleen. Moreover, the presence of eIF5A^Hyp^ co-expressing cells was not enhanced in diseased tissue; however, larger cohorts are required to quantitate the presence of these cells and definitively determine correlation with disease.

Our findings in the pancreas demonstrate that eIF5A^Hyp^ is expressed in both the exocrine and endocrine compartments in mouse and human. Strikingly, the immunostaining analysis revealed that the PP cell population exhibited the most robust immunostaining for eIF5A^Hyp^. Given the over-representation of PP cells in the uncinate region of the pancreas [17], we specifically analyzed tissue sections that contained the uncinate region and found that, regardless of location, PP cells co-expressed eIF5A^Hyp^. Despite evidence that PP cells have a critical secretory function in the brain-gut axis [18] and may serve as a regulator of intra-islet secretion [19], the role of PP cells in the context of diabetes has received little attention. From a developmental perspective, PP cells are predominantly derived from the ghrelin-expressing cell lineage found in the embryonic pancreas [20]; however, the function of eIF5A^Hyp^ in the PP cell population postnatally or any function for eIF5A^Hyp^ in the development of PP cells has yet to be elucidated. Interestingly, expression analysis of 12-lipoxygenase, a factor known to promote inflammation in the setting of diabetes, is also increased in the PP-expressing cell population in pancreas tissue from human donors (collected through nPOD; [21]). Clearly, a greater understanding is required for the role of PP cells in the pathogenesis of diabetes.

Given that much of the published and ongoing work on hypusine biosynthesis in mice has studied eIF5A^Hyp^ in the context of diabetes, we had hypothesized that eIF5A^Hyp^ expression would be identified predominantly in the insulin-producing beta cell population. Our western blot analysis did reveal eIF5A^Hyp^ expression in human islets. Moreover, we observed eIF5A^Hyp^ expression in a purified population of beta cells (tomato+) from mouse islets. Interestingly, the quantitative nature of western blots indicates that the expression of eIF5A^Hyp^ must be lower in the purified beta cells compared with non-beta cells given that PP cells are only a small portion of the tomato(-) non-beta cell fraction whereas the tomato(+) fraction is composed exclusively of beta cells, and we see near equivalent expression of eIF5A^Hyp^ in both sorted populations. This finding is consistent with the unexpected immunofluorescence data, wherein we identified robust expression of eIF5A^Hyp^ in PP-expressing cells. Therefore, together these data support the conclusion that eIF5A^Hyp^ is expressed at a lower level in beta cells compared with the PP cell population.

Our previous finding that pharmacological inhibition of eIF5A hypusination (using the drug GC7) in non-obese diabetic (NOD) mice improved glucose tolerance and preserved beta cell mass [12]. These improvements were also accompanied by reductions in insulitis, which led us to question whether the improvements in beta cell function were due to a direct effect of DHPS inhibition in beta cells, or an indirect effect related to DHPS inhibition in infiltrating immune cells. Our work and that of others suggest a role for eIF5A^Hyp^ and DHPS in promoting T cell and B cell proliferation [12,22,23], which was the basis for our hypothesis that perhaps eIF5A^Hyp^ is differentially expressed in immune cells in individuals with diabetes compared with controls. However, identical expression patterns and abundance was noted in all spleen tissue evaluated. Given our recent findings that deletion of *Dhps* in adult mouse beta cells impacts diet-induced beta cell proliferation (Lavasseur E. et al., *in submission*), we are now investigating eIF5A^Hyp^ expression in other proliferating cell populations as well as diabetes-induced inflammation.

## MATERIALS AND METHODS

### Pancreas and isolated islet cells from mouse

All mice were purchased from the Jackson Laboratory and maintained under a protocol approved by the Indiana University School of Medicine Institutional Animal Care and Use Committee. Total pancreas and isolated islets from wildtype C57BL/6 mice as well as acinar tissue and isolated islets from human donors (Table 3 details human donors) were subjected to immunoblot analysis as previously described [24]. Rabbit anti-eIF5A^Hyp^ ([11]; 1:1000), mouse anti-eIF5A (BD Biosciences; 1:2000) and guinea pig anti-insulin (DAKO; 1:5000) antibodies were used to confirm protein expression in the pancreas.

**Table 3.** Human donor information for islet and acinar tissue preparations.

|  | **Islet Preparation** | **Acinar Preparation** |
| --- | --- | --- |
| Unique identifier | SAMN11578698 | UNOS AGDB487 |
| Donor Age (years) | 57.0 | 24 |
| Donor Sex (M/F) | M | M |
| Donor BMI (kg/m<sup>2</sup>) | 25.8 | 31.7 |
| Donor HbA1c | 5.7 | not tested |
| Origin/source of islets | IIDP<br>(Integrated Islet Distribution Program) | CORE<br>(Center for Organ Recovery and Education) |
| Islet isolation center | The Scharp-Lacy Research Institute | Pittsburgh (AHN) |
| Donor history of diabetes? Yes/No | No | No |
| **If Yes, complete the next two lines if this information is available** |  |  |
| Diabetes duration (years) | Not Applicable | Not Applicable |
| Glucose-lowering therapy at time of death | Not Applicable | Not Applicable |

Mice containing the *RIP-cre* allele (B6.CG-Tg(Ins2-cre)25Mgn/J) [25] were mated with those containing the *R26R*^*Tomato*^ allele (B6.Cg-Gt(ROSA)26Sor^tm14(CAG-tdTomato)Hze/J^) [26] to produce double transgenic animals wherein all the pancreatic insulin-producing beta cells in the pancreas expressed a fluorescent (Tomato) reporter. Pancreatic islets were isolated from *RIP-cre;R26R*^*Tomato*^ mice and *R26R*^*Tomato*^ mice as previously described [27]. The isolated islets from all mice were pooled together and processed for fluorescence activated cell sorting (FACS), which facilitated the separation of islet cells into two populations: tomato-positive beta cells and tomato-negative non-beta cells (islet cells expressing glucagon, somatostatin, ghrelin, and pancreatic polypeptide). Pooled islets were washed with sterile PBS (Fisher Scientific) and incubated in Accutase cell detachment solution (Sigma) for 10 minutes at 37C with constant mixing (1000 rpm). Islet cells were removed from the Accutase solution by centrifugation (500 × g for 1 min) and resuspended in cold buffer containing 2% BSA,1uM EDTA, and equal parts PBS and HBSS (Fisher Scientific). The cells were filtered, collected and incubated with APC viability dye (Zombie NIR-IR dye; BioLegend) per the manufacturer’s recommended protocol. Single-cell suspensions from *RIP-cre;R26R*^*Tomato*^ mice and *R26R*^*Tomato*^ were then sorted using an iCyt Reflection with 100 μm nozzle at 23 psi. Dead cells (NIR-IR+) were excluded; tomato(+) cells and tomato(-) cells were collected into tubes containing sort buffer. Data were analyzed using FlowJo software (Tree Star). Lysates from the two populations of cells were subjected to immunoblot analysis. Rabbit anti-eIF5A^Hyp^ and mouse anti-eIF5A antibodies were used as above, to evaluate the abundance of eIF5A^Hyp^ in the beta cell and non-beta cell populations. Rabbit anti-Pdx1 (Chemicon; 1:1000) antibody was used to evaluate enrichment of the beta cell (tomato+) population.

### Mouse pancreas tissue and immunofluorescence analysis

Pancreas tissue was harvested from wildtype C57BL/6 mice and fixed in 4% paraformaldehyde (Fisher Scientific), cryo-preserved using 30% sucrose, embedded in OCT (Fisher Scientific) and sectioned onto glass microscope slides. Methods previously described for pancreas preservation and immunofluorescence were followed [28]. Pancreas tissue sections (8 um) were stained using the following primary antibodies: guinea pig anti-insulin (DAKO; 1:500), goat anti-pancreatic polypeptide (abcam; 1:200), rabbit anti-eIF5A^Hyp^ ([11]; 1:1000). Secondary antibodies including Alexa-488, Cy3, or Alexa-647 (Jackson Immunoresearch) were used, followed by DAPI (Sigma) to visualize nuclei. Images were acquired with a Zeiss 710 confocal microscope.

### Human pancreas and spleen tissue

Paraffin-embedded tissue sections were obtained from the nPOD consortium (www.jdrfnpod.org). A total of 10 nondiabetic donors, 4 donors with T2D, and 12 donors with T1D (6 autoantibody positive, 6 autoantibody negative) were included in this study (Table 1 and Table 2). Information regarding donors’ demography, histology, and disease status were provided by nPOD. The status of autoantibodies was also determined by nPOD as previously described [29].

### Immunofluorescence analysis of human tissues

Immunofluorescent staining was performed as previously published [28] with modifications to account for the use of paraffin embedded tissue. Briefly, tissue sections were deparaffinized through graded ethanols (100%, 95%, 85%, 75%, 50%; Fisher Scientific) and then blocked using normal donkey serum (Sigma). Primary antibodies used included guinea pig anti-insulin (DAKO; 1:500), mouse anti-glucagon (Abcam; 1:500), rat anti-somatostatin (abcam; 1:200), goat anti-pancreatic polypeptide (abcam; 1:200), goat anti-ghrelin (Santa Cruz; 1:500), mouse anti-Pax5 (DAKO; 1:200), mouse anti-CD8 (Thermo Fisher; 1:500), mouse anti-CD4 (Leica; 1:500). Secondary antibodies including Alexa-488, Cy3, or Alexa-647 (Jackson Immunoresearch) were used to visualize primary antibodies. DAPI (Sigma) was used to visualize nuclei. Images were acquired with a Zeiss 710 confocal microscope.

## ACKNOWLEDGEMENTS

The authors wish to thank Dr. David Morris and the Flow Cytometry Core Facility at Indiana University School of Medicine for assistance with FACS. Human pancreatic islets were provided by the NIDDK-funded Integrated Islet Distribution Program (IIDP) at City of Hope, NIH Grant # 2UC4DK098085. Human donor acinar tissue was provided by Dr. Rita Bottino at the Center for Organ Recovery and Education (CORE), Pittsburg PA. This work was supported by a JDRF Career Development Award (5-CDA-2016-194-A-N) to TLM, NIH R01 DK60581 to RGM, and a JDRF award to RGM. This research was also performed with the support of the Network for Pancreatic Organ donors with Diabetes (nPOD; RRID:SCR_014641), a collaborative type 1 diabetes research project sponsored by JDRF (nPOD: 5-SRA-2018-557-Q-R) and The Leona M. & Harry B. Helmsley Charitable Trust (Grant#2018PG-T1D053). Organ Procurement Organizations (OPO) partnering with nPOD to provide research resources are listed at http://www.jdrfnpod.org//for-partners/npod-partners/.

## AUTHOR CONTRIBUTIONS

TLM, RGM designed research; TLM, SCC, LRP performed research; TLM, SCC, LRP, RGM analyzed data; TLM, RGM wrote the manuscript; all authors edited and approved the final draft of the manuscript.

## THE PAPER EXPLAINED

### Problem

Previous studies in mouse models of type 1 and type 2 diabetes showed that reducing the hypusinated form eIF5A (eIF5A^Hyp^) resulted in improved glucose tolerance and preserved or enhanced beta cell mass. These findings suggests that eIF5A^Hyp^ may play a critical role in the pathogenesis of diabetes by acting directly on the beta cells and/or altering the adaptive or innate immune responses. Examining tissues from individuals with and without type 1 and type 2 diabetes to determine the expression pattern of eIF5A^Hyp^ and stratification with disease will provide insight into the relevance of eIF5A^Hyp^ in the human disease setting.

### Results

As expected, we identified expression of eIF5A^Hyp^ in the beta cells and exocrine cells in the pancreas as well as immune cells in the spleen; however, there was an unexpected cell-specific enrichment of eIF5A^Hyp^ in the pancreatic polypeptide-expressing PP cells in the pancreas. Moreover, the presence of eIF5A^Hyp^ co-expressing PP cells was not apparently altered with disease.

### Impact

Our data highlights new aspects of eIF5A biology and identified robust expression of eIF5A^Hyp^ in the PP cells of the pancreas.

